# Self-Supervised Electrocardiograph De-noising

**DOI:** 10.1101/2024.01.01.573833

**Authors:** Xiaobo Liu, Jiadong Yan, Yisen Huang, Yubin Wang, Chanchan Lin, Yingxuan Huang, Xiaoqiang Liu

**Affiliations:** First Hospital of Quanzhou Affiliated to Fujian Medical University, Quanzhou, Fujian, China; McConnell Brain Imaging Centre, Montreal Neurological Institute, McGill University, Montreal, Quebec, Canada

**Keywords:** ECG, De-noising, Self-supervision

## Abstract

The electrocardiogram (ECG) records heart-beats and is potentially life-saving. However, the ECG signals (e.g., recorded by the standard ECG monitoring system or the Holter ECG monitoring system) heavily suffered from the noises. Thus, the recorded signals involve meaningful cardiac deflections, other biological waves (e.g., caused by the muscle or electrode movements), and even some noises from the monitoring devices (e.g., the power cables). These noise sources would result in inaccurate analyses of cardiac diseases, thus requiring the ECG de-noising methods in data pre-processing phase before the diagnoses. Previous work provided various ECG de-noising approaches, typically based on some filter algorithms or some wave decomposition algorithms. Most of these approaches did not profoundly consider the ECG signals’ specific data structure, and were not adaptive to the signals recorded by various devices or different skin electrodes. Inspired by the ECG recording theory, we find it available to extract noise information from noisy ECG signals directly. We propose a new ECG de-noising method implemented by the neural network, which de-noise the ECG signals without the supervision of the clean signals. The procedure in the self-supervision is straightforward: we first estimate and simulate the noise signals according to the given noisy ECG signals, and then “subtract” the simulated noises to obtain the de-noised ECG signals by the neural network. Experiments on a public dataset verify that our approach is adaptive to ECG signals from different patients and devices. Also, it is proven that the classification on the ECG signals de-noised by the proposed de-noising methods outperforms those with the traditional de-noising methods.

## I. INTRODUCTION

**E**LECTROCARDIOGRAPH is a non-invasive tool in recording heartbeat signals, which is helpful to the diagnoses of the cardiac abnormalities and disorders. A piece of electrocardiogram (ECG) signal often represents several cardiac cycles recorded by skin electrodes, showing the cardiac activities over time. The cardiac abnormalities can be recognized from the ECGs by the physicians or automatic diagnosis systems [1]–[3]. However, the property of the non-invasion is a double-edged sword, which also makes the recorded signals contain not only useful cardiac deflection signals but also other signals that we regard as noises. These noises are divided into cardiac and extra cardiac [4], which are caused by the patients’ characters and the ECG monitoring devices, respectively. In practice, the ECG monitoring approaches are various, including the standard ECG lead system with 8 or 12 skin electrodes, the dynamic ECG monitoring with one or few skin electrodes, and the Frank system, etc. These approaches followed different monitoring rules and electrode positioning principles, thus causing noises with different patterns.

Besides, in previous study [4], [5], the noises in ECG recording are further summarized into five types, including the power line interface of the devices, the baseline wanders, the electromyogram (EMG) or muscle artifacts, the motion artifacts, and the other instrumental and electrosurgical noises. Further, Singh et al. pointed out [6] that the ECG signals contained channel noises that affected the frequency of ECG signals. These noises contaminating the ECG signals are of different patterns and are varied with patients, devices, doctors. There were many approaches proposed for ECG de-noising based on fuzzy methods [7], [8], various filters [9]–[11], neural networks [12], [13]. These methods, however, were not so adaptive as they were built with some strong priors or assumptions, or were highly influenced by some hyperparameters (e.g., some thresholding-based methods). Thus, the de-nosing methods could not be suitable for all types of ECG signals with noises from different distributions.

A typical ECG signal comprises several cardiac cycles, and each cardiac cycle sequentially consists of a P wave, a QRS complex, a T wave, and some intervals between these waves. Also, there is an interval at the end of each cardiac cycle (we call it “TP-interval”). We regard the signals in the TP-interval as a representation of the noise signals, because cardiac activities in the duration are off. In other words, in the duration of a heartbeat, the signal is the addition of the noise signals and the cardiac signals; in the TP-interval, only the noise signal is recorded because of the cardiac inactivity. However, even if we obtain the distribution of the noises, it is still not obvious how we can remove the noises from the ECG signals, because the noises sampled from a certain distribution can be entirely independent.

Currently, there were some progresses in neural network based de-nosing methods [14]–[17]. Previous work verified that neural networks could restore the corrupted images by extracting the underlying image priors. Further, motivated by the adversarial learning, Lehtinen et al. [14] proposed a method to restore the images with only the given noisy image pairs and obtained considerable performances. These approaches used a collection of corrupted versions of one image to estimate the image prior, where the clean image is regarded as the expectation of the images with various corruptions. Also, a self-supervised de-noising method was proposed [16], using a blind-spot mask rather than the image pairs as in [14]. These approaches provided some solutions to deal with noisy data without a clean version. Further, a generative adversarial network made use of the hierarchical noises [18] to facilitate generate the detailed image patterns, which verified that the neural network could extract the clean signals from the noise-disturbed versions.

In this paper, we borrow the ideas of self-supervised image de-noising methods for ECG signal de-noising. An essential improvement is that we treat the TP interval in each cardiac cycle as a sample of the noise, and directly extract the information of ECG noise by computing the statistical parameters of the noises. Based on the statistical parameters, we can simulate a noise of the same pattern and manage to “subtract” the noises from the raw ECG signals by using a neural network. Following the insight, we use a U-shape auto-encoder neural network to deal with the ECG signals. In processing, the noisy ECG signals and the simulated noises are fed into the proposed auto-encoder, which predicts the corresponding clean ECG signals. Experiments on the tianchi ECG dataset^1^ verify that our ECG de-noising method is helpful, and a deep learning classifier obtains better performances with the denoised ECG signals by our proposed method. There are three major contributions in this work:

- We propose a neural network based method for ECG signal de-noising, which is an adaptive model can denoise the ECG signals without the supervision of the clean signal.
- We make use of the property of the TP interval in the ECG signals by regarding the signals in the TP interval of a sample of the noise, which helps the deep learning model learn to “substract” the noises from the ECG signals.
- Experiments show our proposed method benefits to disease diagnosis (classification) based on the ECG signals, which further verify that the proposed approach works well in ECG de-noising and remains clinically useful information.

## II. Preliminaries

### A. Backgrounds

Electrocardiography is a non-invasive tool for heartbeat signal recording, which is widely used in diagnosing cardiac abnormalities and functional disorders. Typically, a piece of ECG signal is composed of several cardiac cycles (full heartbeats), and each cardiac cycle consists of a series of deflections, as shown in Fig. 1. The major deflections in a normal cardiac cycle are P wave, QRS complex, and T wave, determined by several turning points: P, Q, R, S, and T. Also, there is a small deflection, called the U wave, following the T wave. But, since the U waves are often tiny, it is often hid by the ECG noises. A cardiac cycle is defined by the period from the beginning of the atria contraction to the end of the ventricular relaxation, and there are no cardiac activities recorded in the period of the TP interval, thus we regard the recorded signals as the sample of noises. Unlike the de-noising tasks on other data, there is a short-cut for ECG de-noising because the signals in the TP intervals can be treated as representations of noises.

**Fig. 1.**
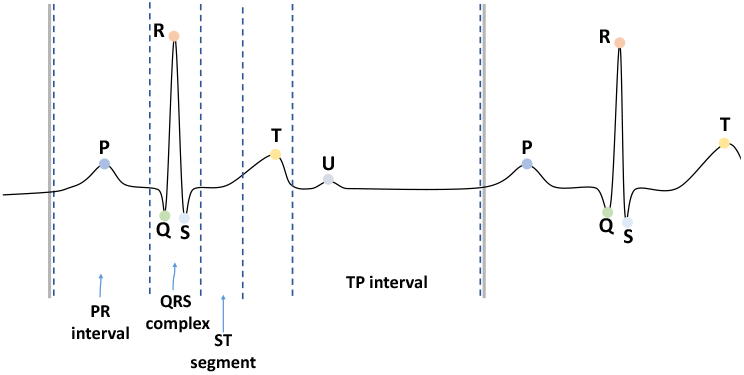
Illustrating the structure of a normal ECG signal. Note that the U wave is very tiny and is often hidden by the noises or other signal components.

### B. ECG signal de-noising algorithms

Due to the clinical importance of ECG monitoring, there were various ECG de-noising methods proposed in the previous work. Various filtering algorithms were proposed earlier, such as the adaptive filter [19], low-pass filter [20], filter bank-based method [21], and the Kalman filter [22]. Later, some non-linear filtering algorithms (e.g., the Bayesian filtering [23] and the non-linear projective filtering [24]) and some variants of the least square algorithms [25], [26] were also introduced for ECG de-noising tasks. The fuzzy methods [7], [8] have also been employed in ECG de-noising researches.

However, the most popular algorithms followed the signal decomposition techniques in the time domain [27]–[30], especially based on the discrete wavelet transform (DWT) algorithm and the empirical mode decomposition (EMD). The traditional de-noising method, non-local means (NLM) [31], [32], was proven to be performance-beneficial in the ECG de-noising task, and we attributed it to the full use of the global information and the pattern comparison [6], [33], [34]. Interestingly, the progressive image de-noising (PID) [35] was used in removing the ECG signals from the electromyogram [36], where the amplitude of the electromyogram is modeled as white Gaussian noise.

With the developments of neural networks, some neural network based methods were explored for ECG de-noising [37]– [39]. In [37], [39], the auto-encoder neural networks were used in ECG de-noising, which achieved better performances than most traditional ECG de-noising methods. A generative adversarial network (GAN) for ECG de-noising model [38] was proposed, which was claimed to be the first research to use a GAN for the ECG de-noising purpose. This work verified a neural network based ECG de-noising method could be adaptive to multiple noise types, and the neural network based ECG de-noising method is promising. The drawback is that these methods required clean ECG signals and noisy ECG signals in training, which is unpractical because the clean ECG signals are hard to obtain in practice.

### C. Neural network based de-nosing algorithms

There were many neural networks proposed for data de-noising. This field’s beginnings are CSF [40] and TNRD [41], which outperformed the state-of-the-art traditional methods, Block Matching 3D (BM3D) [42], with light neural networks. Zhang et al. [43] used the residual learning manner [44] and batch normalization [45] in the de-noising model and obtained promising performances. Xu et al. [46] proposed a neural network for the problem that the image channels are contaminated with different noises. These work verified the neural networks to be potential in de-noising, but they only explored the data with the noises from the same patterns and needed the clean data in training. However, it is unreasonable to train a mapping to transform the noisy ECG signals to the clean signals in supervision, as the clean data is actually inaccessible.

Recently, some researchers explored the self-supervised de-noising methods under the conditions that the clean data is absent. These researches were more close to the real scene. A good beginning is that Dmitry et al. [15] verified the neural network could learn the image prior without direct supervision. Then, some researchers found ways to de-noise ECG signals with only the noisy data [14], [16], [17]. In [14], the de-noising problem is reformulated into the problem of empirical risk minimization, and thus obtained better performances than those methods with clean data. But this method required image pairs with different noises added on the same images. Further, a specific blind-spot receptive field was proposed to predict the pixel values by the nearby pixel values except for itself, which did not require the image pairs [16]. A similar idea was explored in [17], de-noising data without data pairs. These work above showed that neural networks could extract the underlying data prior, and the de-noising neural network can be trained in the self-supervision manner, without the supervision of the clean data.

## III. Methods

### A. Insight

In this paper, we introduce a self-supervised method for ECG de-noising. From the view of the signal addition, we define a noisy ECG signal *x* by:

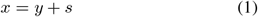

where *y* is the clean ECG signal and *s* is the noise signal. Most of the known work managed to extract the noises from the raw ECG signals without considering the special data structure of the ECG signals. That is, the TP interval in an ECG represents a period of electrical inactivity, *y*_TP_ = 0, and we can obtain:

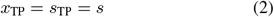

In the other word, we suppose that the noise in an ECG signal can be observed in the TP interval (i.e., *s*_TP_ = *s*), and thus we can model the noises by analysing the TP interval. In the following, we would show how we perform this insight and propose an ECG de-noising neural network for the ECG noise removal.

It is naive to obtain the clean ECG signal *y* by removing the noises as *y* = *x* − *s* in theory, but is complicate in practice. A typical neural network trained with a large number of input-target pair (*x*_*i*_, *y*_*i*_) in supervised learning strategy follows the empirical risk minimization principle to learn *f*_*θ*_(*x*) = *y*, by:

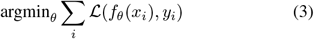

where *f*_*θ*_ indicates a neural network specified by the loss function *L*. Assume that we have a set of noisy ECG signals {*x*_1_, *x*_2_, …, *x*_*i*_, … } of one clean ECG signal, where the elements can be defined by:

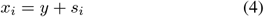

A typical statistical strategy to obtain the clean signal is to find a *y*′ with the smallest average deviation under the measurements determined by a loss function ℒ, as:

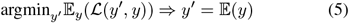

Similar to the analyses in [14], *y*′ can be the except or the median of the noisy signals, with different loss functions. For input-target pairs with noises, the empirical risk minimization principle can be reformulated by:

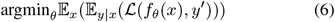

In Eq.(6), we desire *f*_*θ*_(*x*) converge to *y*′ instead of *y*, which is according to the fact that a noisy data actually have many versions of clean data, as discussed in [14]. Eq.(6) suggests that, if an estimation of the noise is obtained, it is available to de-noise data without clean data, since:

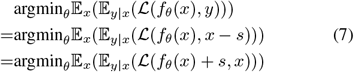

Motivated by Eq.(7) and the literature [14], we design the model to learn

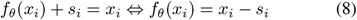

That means, if we can obtain the noise *s*_*i*_ for the recorded ECG signal *x*_*i*_, it is available to predict the clean ECG signal by “substract” the noise, as

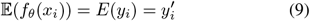

Due to the specific structure of an ECG signal mentioned in Sec. II, it is available to simulate noise signals identically distributed with the recorded ECG signals’ noises. The TP interval is a period where the electrical is inactive and thus only contains the noise signals. A naive method is to regard the noise as the white Gaussian noise and compute the distribution parameters. A more elaborate method can employ a GAN [47] to synthesize noise signals according to a trained neural discriminator. In the previous work, ECG noises were often regarded as white Gaussian noise. Here we follow the white Gaussian noise assumption in this work, and compute the parameter (the expectation *μ* and standard deviation *σ*) of the Gaussian distribution for noises, and synthesize noise signals based on the computed parameters.

Suppose we have obtained a collection of the synthesized noises {*s*_1_, *s*_2_, …, *s*_*n*_} (*n* is a very large number) by a certain noise generator as mentioned above, in which the elements are independently and identically distributed with the noise *s* in the ECG signals. We can build a neural network for signal mapping (*f*_*θ*_(*x*)), and employ a regression loss function to guide the regression of Eq. (8) without the supervision of the clean signal *y*.

### B. Segmentation for the cardiac cycles and TP intervals

To obtain the noise sample of an ECG signal, we should first segment the TP intervals and the cardiac cycles from the ECG signals. In this work we process every cardiac cycle signal separately. We prepare the samples containing the cardiac cycles and the corresponding TP intervals as (*x*_*i*_, *s*_TP_). Previous work often segmented an ECG signal into 7 segments, including P wave, PR-segment, QRS complex, ST-segment, T wave, TP interval, but we only need to segment the TP interval and divide a piece of ECG signal into several cardiac cycles. We train a 1D U-Net [48] to segment the TP intervals and the cardiac cycles. In practice, as the beginnings and ends of the TP interval and the cardiac cycles are distinguishable, we only need to annotate around 100 ECG signal samples for training the 1D U-Net, which is very cheap. After training, the 1D U-Net is then performed on the samples of the whole dataset. Then, a noise generator synthesizes the noises identically distributed with the original noises, in which we compute the expectation *μ* and standard deviation *σ* under the white Gaussian noise assumption. As there might be a U wave in the TP interval, we compute *μ* and *σ* from the second half part signals in the TP interval in the view of accuracy.

### C. The proposed ECG de-noising model

Some known neural networks for ECG de-noising were built following the architecture of the auto-encoders [37], [39], and achieved state-of-the-art performances. Here we build an auto-encoder neural network for ECG de-noising, as shown in Fig. 2 (b). The input noisy ECG signals *x* are processed to obtain a low-dimensional hidden state *z* by the encoder part of the neural network, and the hidden state is used to restore the ECG signal by a reverse model (decoder). As shown in Fig. 2, the encoder part contains several 1D convolutions and down-sampling operations (pooling), as well as several 1D residual basic blocks [44]. The residual basic blocks are the 1D version of the residual block in the ResNet [44]. In contrast, the decoder part consists of several 1D convolutions and up-sampling operations (interpolation). Following the analysis in Sec. III-A, we denote the output of the de-noising neural network as *f*_*θ*_(*x*) = *x*′, and then we add one of the synthesized noise signals onto the predicted signal, by *x*′ + *s*_*j*_, *s*_*j*_ ∈{*s*_*j*_ *s*_1_, *s*_2_, …, *s*_*n*_ }. As the synthesized noises are with an identical distribution to the original noise, we randomly sample one of the noises from the synthesized noise set in every training iteration, and train the model to learn the equivalence as in Eq.(8). In practice, we follow the white Gaussian noise assumption to synthesize {*s*_1_, *s*_2_, …, *s*_*n*_}, and add a sampled noise directly onto the predicted signals. A loss function, here we set L the *L*_2_ loss function as default, by:

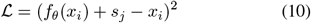

where *s*_*j*_ is sampled in every iteration by the no replacement sampling strategy.

**Fig. 2.**
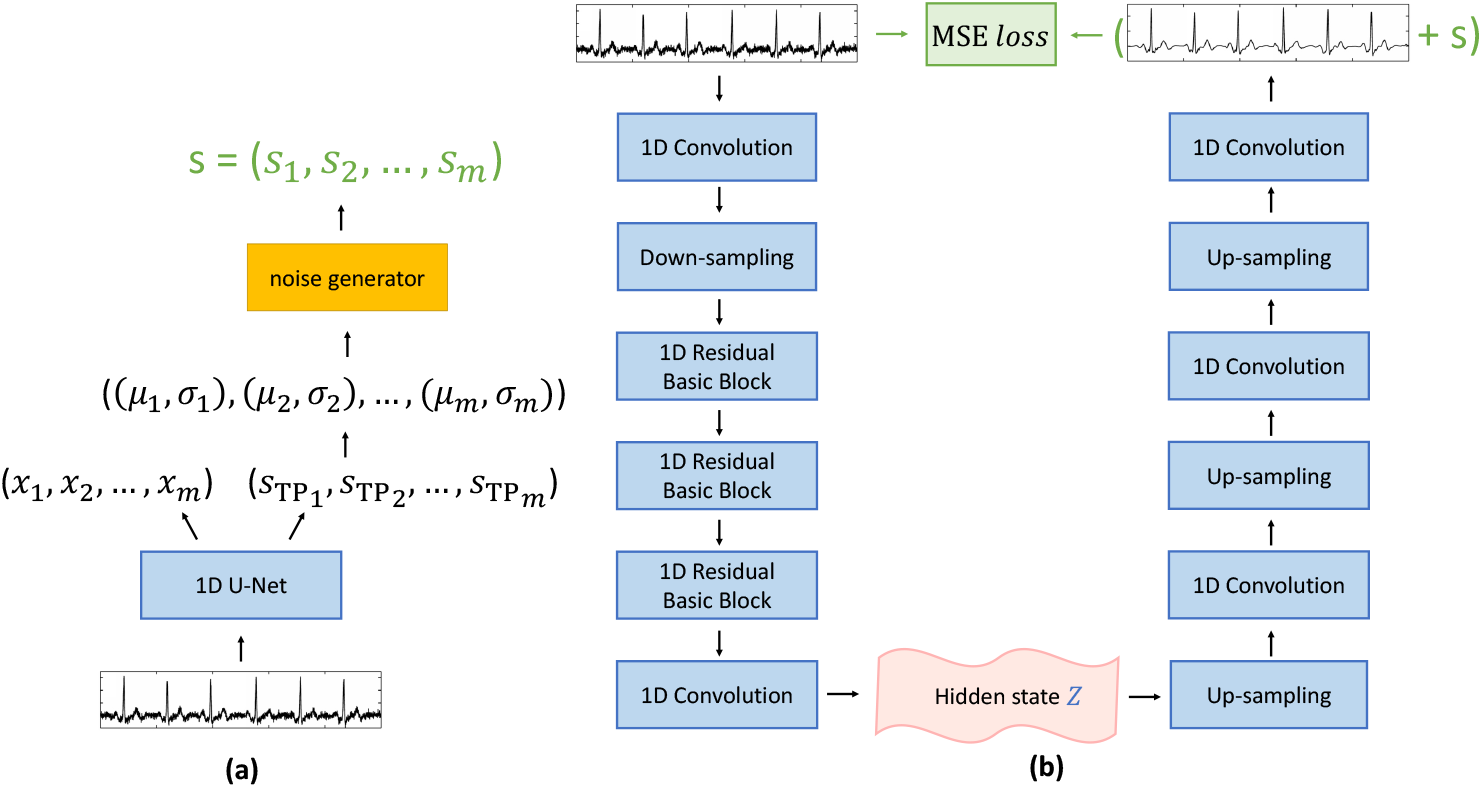
An illustration of the ECG de-noising system. (a) illustrates the procedure of the noise synthesis with 1D U-Net for TP interval segmentation and the cardiac cycle division. The noise generator is discussed in Sec. III-A. (b) illustrates the architecture of the de-noising auto-encoder neural network. Note that the noise “s” in Subfig. (b) is synthesized by the noise generator in Subfig. (a).

### D. Training and inference procedure

The ECG de-noising in our work operates in two discrete steps. First, we annotate several TP interval range on some ECG signal samples, and train a 1D U-Net for the automatic TP interval segmentation. After training the 1D U-Net, the trained U-Net is run on the ECG signals in the training set and test set, and obtain the TP intervals in every cardiac cycle in ECG signals. The TP interval is regarded as the complete noise signal, and a noise generator is used to synthesize noises of the same distribution. There are many ways to build a noise generator, and we use a naive solution by using the statistical method, as discussed in Sec. III-A under the white Gaussian noises assumption. A proposed 1D auto-encoder neural network is used to predict the clean ECG signal, and the noises from the noise generator are added to the predicted clean ECG signals. The training of the 1D auto-encoder neural network is under the guidance of the MSE loss function, as shown in Eq.(10).

In the inference, the ECG signals are first processed by the1D U-Net to obtain the cardiac cycles *x*. Then, the trained auto-encoder deals with the cardiac cycle signals *x* to predict the clean ECG signal, by *y*′ = *f*_*θ*_(*x*).

## IV. Experiments

### A. Data Preparation

To evaluate our proposed method for ECG signal de-noising, we conduct experiments on a selected ECG dataset from the Alibaba Tianchi Cloud Competition dataset^2^ containing 18,604 samples, where most of the ECG signals represent the characters of one or few heart diseases. The ECG signals recorded at a frequency of 500 Hertz with the length of 10 seconds, which contains 8 most common heart disease categories. We randomly divided the selected dataset for training and testing by 8:2, as the official test set is not publicly provided. The Tianchi ECG dataset do not provide the cardiac cycle segmentation ground truth for cardiac cycles, so we annotate the cardiac cycle divisions and the TP interval divisions for around 100 samples for training the 1D U-Net. To directly evaluate the de-noising effects, we first de-noised the ECG signals by using the package [49], and regard the de-noised signals as the clean data. Then we add the Gaussian noises onto the clean data to synthesize the noisy ECG signals, and thus we have the ground truth in evaluating the de-noising performances. To evaluate the de-noising effects on the raw ECG signals, we compare the classification performances on raw ECG signals de-noised by our proposed methods and other de-noising methods. In the experiments, we only used the II lead signals. We first normalize the ECG signals by the linear scaling normalization to 0 ∼ 1, as:

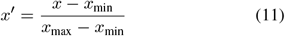

where *x*_min_ and *x*_max_ indicate the minimum and the maximum values in the ECG signals.

### B. Experimental Setup

We use PyTorch 1.7 on Python 3.8 on RTX2080Ti GPUs to implement our proposed methods and the auto-encoder. In training, we train the auto-encoder with batch size 256 in 150 epochs. The learning rate is set 0.1, and is reduced to 0.01 at the 50-th epoch and is reduced to 0.01 at the 150-th epoch. We use the SGD optimizer with momentum 0.9 [52]. We evaluate the de-noising effects via the indicator Peak Signal-to-Noise Ratio (PSNR) and the structural similarity index (SSIM index). Formally, the PSNR of the predicted clean signal (*p*) is defined by:

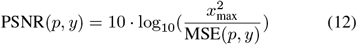

where 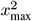 indicates the maximum value on the signal, and the MSE(*p, y*) computes the mean-square error between the clean signal ground truth and the predicted clean signal. The SSIM index is computed by:

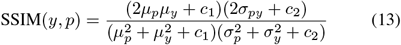

where *c*_1_ and *c*_2_ are two constants. The SSIM index was first proposed for image quality evaluation [53], and here we used a 1D version for ECG signals.

### C. De-noising effects on artificial noises

In this section, we evaluate the de-noising effects of various de-noising methods (including ours, the comprehensive method in the NeuroKit2 package [49], the Band-pass filter [50], the Butterworth filter [51], and the Chebyshev filter [51]) on the ECG signals with artificial noises. The artificial noises are sampled from the Gaussian distribution, with different standard deviations. As shown in Table I, it is obvious that our method outperforms the previous methods on PSNR and SSIM. With bigger noises, the performances of all the methods decrease to some extent, but our method still obtains considerable performances, even with the noise *σ* = 25. To better inspect the effects intuitively, we illustrate the de-noising results in Fig. 3. One might see that our method well de-noise the ECG signals and remain the deflection shapes. In contrast, the other methods can not clearly distinguish the noise signals and the ECG defections with bigger noises (e.g., *σ* = 20 and *σ* = 25), resulting in the deflection distortion.

**TABLE I.** The performances of the de-noising methods. The best performances are marked **in bold**.

| Method | $\sigma=5$ | | $\sigma=10$ | | $\sigma=15$ | | $\sigma=20$ | | $\sigma=25$ | |
| --- | --- | --- | --- | --- | --- | --- | --- | --- | --- | --- |
|  | PSNR | SSIM | PSNR | SSIM | PSNR | SSIM | PSNR | SSIM | PSNR | SSIM |
| Ours | **31.33** | **0.9255** | **27.71** | **0.8797** | **23.43** | **0.8108** | **21.58** | **0.7582** | **22.40** | **0.7583** |
| NeuroKit2 [49] | 25.52 | 0.8977 | 23.04 | 0.8383 | 21.29 | 0.7544 | 19.48 | 0.7067 | 18.03 | 0.6539 |
| Band-pass filter [50] | 11.92 | 0.5415 | 11.98 | 0.5096 | 11.95 | 0.4609 | 11.93 | 0.4152 | 11.58 | 0.3898 |
| Butterworth filter [51] | 20.32 | 0.7884 | 19.72 | 0.6432 | 18.71 | 0.5211 | 17.53 | 0.3789 | 17.42 | 0.3966 |
| Chebyshev filter [51] | 21.11 | 0.7920 | 20.43 | 0.6883 | 19.56 | 0.5717 | 19.19 | 0.4792 | 18.72 | 0.4590 |

**Fig. 3.**
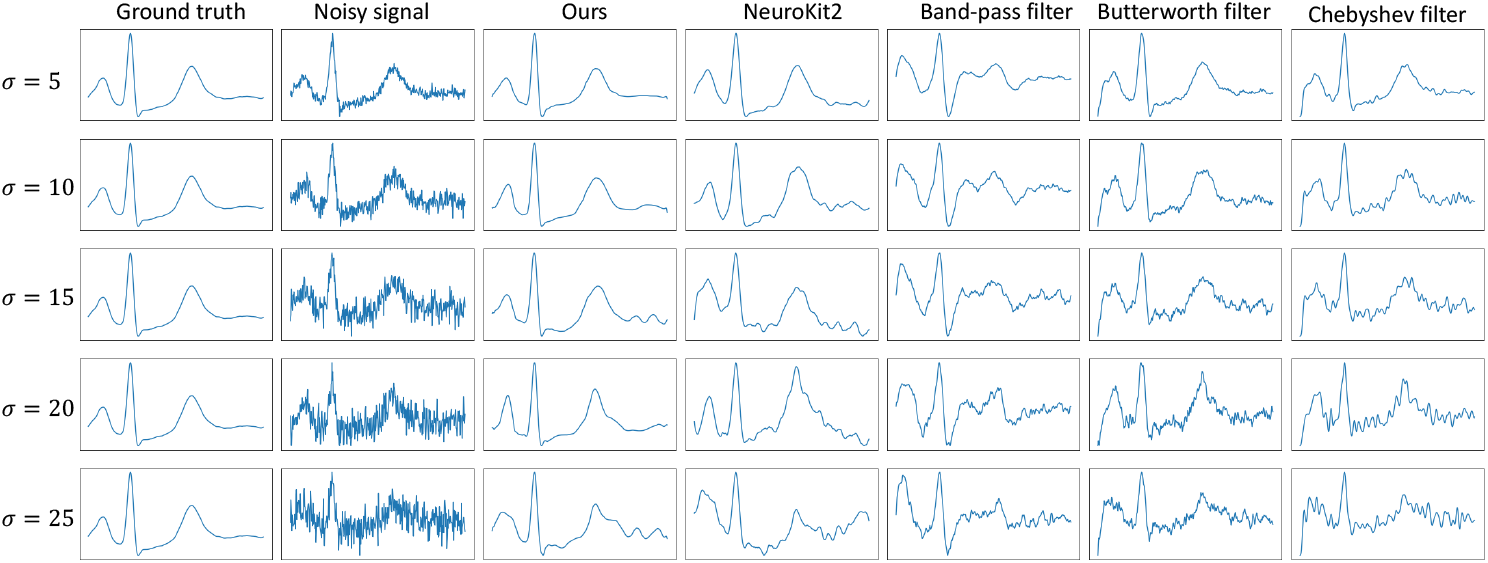
An illustration of the de-noising effects. The noisy ECG signals in the second column are obtained by simulation.

### D. Classification with the de-noised raw ECGs

In the experiments of Sec. IV-C, the noisy ECG signals are obtained by the simulation, which cannot thoroughly verify that our method suits the raw ECG signals. As there are no clean versions of the raw ECG signal, PSNR and SSIM cannot be computed. Thus, we indirectly evaluate the de-noised results by comparing the classification performances on the de-noised ECG signals with an identical classifier. Here we use a 1D ResNet-34 [44] as the classifier, which is trained under the guidance of the binary cross-entropy. As shown in Table. II, the de-noising methods all promote the classification, comparing to the first line. It is evident that the classification performances with the ECG signals de-noised by our method obtain the best performances by clear margins.

**TABLE II.** The classification performances on the ECG signals de-noised by various de-noising methods. The best performances are marked **in bold**.

| De-noised Methods | precision | recall | Micro F1 |
| --- | --- | --- | --- |
| No de-noising | 0.905 | 0.792 | 0.845 |
| Ours | **0.949** | **0.873** | **0.909** |
| NeuroKit2 [49] | 0.934 | 0.841 | 0.885 |
| Band-pass filter [50] | 0.916 | 0.836 | 0.874 |
| Butterworth filter [51] | 0.919 | 0.839 | 0.877 |
| Chebyshev filter [51] | 0.927 | 0.844 | 0.884 |

## V. Conclusion

In this paper, we introduced a novel idea for ECG signal de-noising in a self-supervision manner, following the assumption that the TP intervals represent the recorded noises. We modeled the noises from the TP intervals and “subtract” the noise signals from the ECG signals. In the implementation, an auto-encoder was built to predict the clean signals following our insights, which outperformed the previous de-noising methods on PSNR and SSIM. Also, the classification performances on the ECG signals de-noised by our proposed method were better than those with other de-noising methods. These results suggest that our proposed de-noising method is useful and suit natural noises.

## Acknowledgment

This research is supported by xxx

## Footnotes

1 https://tianchi.aliyun.com/competition/entrance/231754/information?lang=en-us

2 https://tianchi.aliyun.com/competition/entrance/231754/introduction

## Notes

### Competing Interest Statement

The authors have declared no competing interest.

